# Effects of N, P, K on Yield and Quality of Jujube in the Loess Hilly Region and Its Fertilization Model

**DOI:** 10.1101/2020.09.01.277301

**Authors:** Shenglan Ye, Tiancheng Liu, Yulu Wei

## Abstract

The aim of this study is to explore the optimal N, P and K fertilization model suitable for pear-jujube in the mountain of northern Shaanxi in 2016 years. This experiment use 3-factor, saturated optimal design for quadratic fertilization scheme. The effects of different fertilization treatments on the yield and quality of pear-jujube were studied through field experiments. And comprehensive evaluation based on the quality of pear-jujube. The results showed that N_1_P_3_K_3_ has the highest yield, which is 48% higher than the control (CK). The effect of nitrogen, phosphorus, and potassium on yield is potassium fertilizer (positive effect)> phosphate fertilizer (positive effect)> nitrogen fertilizer (negative effect). Phosphate and potassium fertilizers have significant effects on increasing the content of soluble solids. Application of potassium fertilizer alone can significantly increase the content of reduced Vitamins c. The combined application of nitrogen, phosphorus, and potassium reduced the content of reduced Vitamins c. N_3_P_3_K_1_ treatment can significantly increase the total sugar content in fruits. Single application of phosphate and potassium increases the content of organic acids in fruits. Other fertilization treatments have significant effects on reducing the content of organic acids in fruits. The sugar-acid ratio of N_3_P_3_K_1_ is significantly higher than CK, which had an important effect on improving the taste. A high amount of potassium fertilizer has a significant effect on increasing the total flavonoid content in fruits. The interaction of nitrogen and phosphorus will reduce the total flavonoid content. The effect of nitrogen, phosphorus, and potassium on quality is potassium (positive effect)> nitrogen (positive effect)> phosphate (positive effect). Comprehensive analysis, the optimal fertilization amount when the target yield is 23000 ∼ 27000 kg·hm^-2^ and the quality score is above 85 is nitrogen (N) 406.93 ∼ 499.31 kg·hm^-2^, phosphorus (P_2_O_5_) 203.94 ∼ 297.08 kg·hm^-2^, and potassium (K_2_O) 285.47 ∼ 322.82 kg·hm^-2^.

## Introduction

The ecological environment of the Loess Plateau in northern Shaanxi is fragile. The main reason is that vegetation is scarce; soil erosion and land desertification are more serious. The intensity of precipitation and the reclamation of steep slopes have increased soil erosion. As a result, the yield of crops is low and unstable, which seriously restricts the development of the local economy and the improvement of people’s living standards. The analysis by Li Song [1] shows that the development of slope orchards is one of the important ways to improve the ecological environment and economic level of the Loess Plateau. Fertilizer application is an important factor to maintain the quality of the orchard. Formula fertilization can protect the environment and improve fruit yield and quality. Fertilizer input is crucial for sustainable agricultural production that provides sufficient food for the world ^[2]^. And it plays an important role in food safety ^[3]^. The quality of fruit is an important aspect of the value of the commodity. Bie Zhixin et al [4] showed that the combination of nitrogen, phosphorus and farm manure could significantly extend the storage period (post-maturity) of kiwi fruit. The quality and flavor of kiwi fruit have been significantly improved. He Zhongjun et al [5] showed that potassium fertilizer at the same nitrogen and phosphorus level can increase the content of soluble sugar, Vc, hardness, sugar-acid ratio and first-grade fruit rate in kiwifruit, and reduce the content of titratable acid. Wang Qin et al [6] found that adding potassium fertilizer can significantly improve the quality of apples. Zhao Zuoping [7] studied the influence of different fertilizer ratios on apples on this basis. The results show that nitrogen, phosphorus and potassium deficiency treatment will increase the small fruit of Fuji apple. Nitrogen application significantly increased the fiber length and fiber specific strength of hybrid cotton, but the application of nitrogen fertilizer had little effect on it [8-9]. The main quality traits of rape (oil content, erucic acid, arachidonic acid, linolenic acid, glucosinolate, linoleic acid) are significantly correlated with nitrogen application rate [10]. Therefore, a lot of research on fertilizer application has been carried out to improve the ecological environment and economic situation in northern Shaanxi. Existing studies include the application of amino acid fertilizer [11], nitrogen fertilizer [12], and potassium fertilizer [13]; study the effects of combined application of nitrogen, phosphorus and potassium and single application of phosphorus fertilizer on the growth of jujube young trees [14-15]; Nitrogen, phosphorus and potassium absorption and utilization [16]; study the change of nutrients in pear date [17]. However, the output and quality of jujube trees have a great relationship with varieties and soil environment. The theoretical basis of the existing research is not suitable for increasing the yield of pear jujube on the slopes of northern Shaanxi. The combined application of chemical fertilizers will not destroy the soil structure of loessial soil in the northern slope of Shaanxi. It can promote the growth and development of crops and improve the soil physical properties of loessial soil. This plays an important role in improving the ecological environment [18]. To this end, this experiment studied the effects of nitrogen, phosphorus, and potassium fertilizers on the growth, yield, and quality of pear jujube in the drought-irrigated hilly areas of northern Shaanxi Loess Hilly Region, with a view to obtaining a reasonable fertilization program for producing high-yield and high-quality pear jujube. It provides a theoretical basis for improving the low output of pear jujube in the Loess Hilly Region of Northern Shaanxi, developing the pear jujube industry and scientific fertilization.

## Materials and methods

### Overview of the study site

The test was carried out in in Mengzi Village, Mizhi County, Yulin City where is pear jujube planting base. It locates at north latitude 37°43′ ∼ 38°08′and east longitude 100°15′∼110°16′. The landform of test area belongs to the typical hilly and gully region of the Loess Plateau. Soil erosion and land desertification are serious. It is a semi-arid climate in the temperate zone. The climate is dry, the sunlight is sufficient, and the temperature difference between day and night is large, which is a natural resource necessary for the growth of jujube trees. The average annual temperature is 8.5 °C. Sunshine hours are 2716 hours. Frost-free period is 160 ∼ 170 days. The average annual rainfall is 451.6mm. The seasonal distribution of precipitation is uneven. There is little rainfall from April to June, and most of them are invalid precipitation below 10mm. Rainfall from June to September accounts for 74.3% of the whole year, and there are many heavy rains, which makes a considerable part of precipitation form runoff loss. The test soil is mainly Lossiah soil. The soil is relatively barren. The soil bulk density is 1.21g / cm3. The available N, P, and K contents in the soil were 34.73, 2.90, and 101.9 mg / kg, respectively. The soil organic matter content is 2.1g / kg. The pH is 8.6.

### Experimental design

The test tree was 7 years old with uniform growth and good growth. The test site is a horizontal terrace on the slope. The soil water and fertilizer and other factors along the contour line are basically the same. The planting density is 3m × 2m. The fertilization design scheme is shown in Table 1. The test is designed with 10 treatments. Individual plant is one treatment, and a total of 5 replicates are set. Drip irrigation was carried out at the germination stage, flowering stage and fruit setting stage. The total irrigation quota is 135 m^3^/hm^2^. Fertilization method: Dig a ditch 20 cm deep, 40 cm wide and 40 cm long under the canopy and on the outer sides. Fertilizer and soil are mixed 1:1 and spread in the ditch, then covered with soil. On April 26, 2016, all phosphate and potassium and 50% nitrogen fertilizers were applied. On July 21 (fruit expansion period), the remaining 50% nitrogen fertilizer was applied as top dressing. The type of fertilizer used is urea (N content is 46%), superphosphate (P_2_O_5_ content ≥ 12%) and potassium sulfate (K_2_O content ≥50%).

**Table 1.** Three factors D-saturation optimal design of N, P and K

| Treatment | Code |  |  |
| --- | --- | --- | --- |
|  | X<sub>1</sub> | X<sub>2</sub> | X<sub>3</sub> |
| CK | -1 | -1 | -1 |
| N<sub>3</sub> | 1 | -1 | -1 |
| P<sub>3</sub> | -1 | 1 | -1 |
| K<sub>3</sub> | -1 | -1 | 1 |
| P<sub>2</sub>K<sub>2</sub> | -1 | 0.1925 | 0.1925 |
| N<sub>2</sub>K<sub>2</sub> | 0.1925 | -1 | 0.1925 |
| N<sub>2</sub>P<sub>2</sub> | 0.1925 | 0.1925 | -1 |
| N<sub>1</sub>P<sub>3</sub>K<sub>3</sub> | -0.2912 | 1 | 1 |
| N<sub>3</sub>P<sub>1</sub>K<sub>3</sub> | 1 | -0.2912 | 1 |
| N<sub>3</sub>P<sub>3</sub>K<sub>1</sub> | 1 | 1 | -0.2912 |

### Measurement indicators and methods

Yield: After the fruit is mature, the single plant single-receipt method is adopted. Every time the fruit is picked, it is recorded until the fruit picking is completed. Calculate individual yield and convert to total yield.

Jujube fruit size classification: After the jujube fruit is ripe, use a Vernier caliper to measure the longitudinal diameter of the fruit on the secondary branches in different directions. Each tree measures 30 jujube fruits. According to the calculation of the maximum longitudinal diameter and the minimum longitudinal diameter, the graded section is obtained. Fruit diameter less than 32.17mm is common fruit. Fruit diameters between 32.17mm-42.34mm are high-quality fruits, and fruit diameters greater than 42.34mm are exceptional fruits.

Moisture content of fruit: The moisture content of fruit is determined by drying method.

Water content (%) = (fresh weight-dry weight) / fresh weight × 100%

Soluble solids were measured using a 2WAJ-Abbe refractometer. Reducing vitamin c was determined by 2,6-dichloroindophenol titration [19]; the total sugar content was determined by 3,5-dinitrosalicylic acid colorimetric method [20] Organic acids are determined by acid-base titration [19,21], sugar-acid ratio = total sugar/organic acid. The total flavonoids were extracted by ultrasonic wave and then determined by sodium nitrite-aluminum nitrate-sodium hydroxide color method [22-24].

Comprehensive judging standard of jujube quality: Evaluation is based on the general nutrients and medicinal nutrients of jujube. Refer to the evaluation criteria of crop quality such as taro [25] and onion [26]. Six quality indicators of moisture, fruit shape index, soluble solids content, sugar-acid ratio, reduced Vc and total flavonoid content were comprehensively scored in order of importance (out of 100 points). For each index, the best processing value is full marks. The percentage of the measured value of the index in the optimal processing value of the processing is the actual score of the processing index. The sum of the weight values of all the quality index scores of each process is the overall quality score of each process. The total flavonoid content weight is 0.3. The sugar-acid ratio and the reduced Vc weight are both 0.2. The water content, fruit shape index and soluble solids weight are all 0.1.

### Data statistics and analysis

The test data was compiled using Microsoft excel 2010 and SPSS.18.0, and drawing was using MATLAB 7.1.

## Results

### Effects of different fertilization treatments on the output of pear-jujube

The yield of fertilization treatment is significantly higher than control treatment. The output of K_3_, P_2_K_2_ and N_1_P_3_K_3_ is higher than that of other treatments. The yield of N_1_P_3_K_3_ treatment is the highest, reaching 25950.6 kg/hm^2^, 48% higher than the control. It shows that fertilization can increase the yield of jujube trees. Among them, the increase of potassium fertilizer application has a significant effect on increasing yield. Combined application of phosphorus and potassium can further increase the output of red dates on the basis of single application of potassium fertilizer. The reasonable combination of nitrogen, phosphorus and potassium is more beneficial to the increase of jujube yield. (Table 2)

**Table 2.** Effect of N, P and K on the yield of jujube kg·hm^-2^

| kg · hm<sup>-2</sup> |  |  |  |  |
| --- | --- | --- | --- | --- |
| Treatment | Repeat 1 | Repeat 2 | Repeat 3 | Average |
| CK | 17188.45 | 17486.35 | 17822.31 | 17499.04±258.93g |
| N<sub>3</sub> | 20392.92 | 20504.06 | 19904.73 | 20267.24±260.31e |
| P<sub>3</sub> | 18066.81 | 18514.32 | 18641.43 | 18407.52±246.44f |
| K<sub>3</sub> | 22778.55 | 23444.49 | 23239.61 | 23154.22±278.50c |
| P<sub>2</sub>K<sub>2</sub> | 25328.37 | 24456.16 | 25263.87 | 25016.13±396.83b |
| N<sub>2</sub>K<sub>2</sub> | 21958.68 | 22440.42 | 23386.92 | 22595.34±593.28cd |
| N<sub>2</sub>P<sub>2</sub> | 21382.70 | 22123.97 | 22344.97 | 21950.55±411.54de |
| N<sub>1</sub>P<sub>3</sub>K<sub>3</sub> | 25755.73 | 25956.95 | 26139.05 | 25950.58±156.56a |
| N<sub>3</sub>P<sub>1</sub>K<sub>3</sub> | 22668.78 | 22857.22 | 22896.23 | 22807.41±99.31c |
| N<sub>3</sub>P<sub>3</sub>K<sub>1</sub> | 21364.10 | 22311.17 | 20771.43 | 21482.23±634.12e |

Note: Different letters in the same year of the same column indicate that the difference between treatments is 5% significant; the same below.

### Effects of different fertilizer treatments on the quality of pear-jujube

Fertilization can significantly increase the water content of jujube fruit. The N_1_P_3_K_3_ treatment had the highest water content, which was 4.17 percentage points higher than the control. P_2_K_2_ treatment had the highest soluble solids content, which was 2.19 percentage points higher than the control. Phosphate and potassium fertilizers increase the reduced Vc content of jujube fruit and reduce the total sugar content. The total sugar content of P_2_K_2_ treatment was significantly higher than that of phosphate fertilizer and potassium fertilizer. Phosphate fertilizer and potassium fertilizer will increase the organic acid content of jujube fruit. Single application of nitrogen fertilizer or combined application can reduce the content of organic acids. The N_1_P_3_K_3_, N_3_P_1_K_3_ and N_3_P_3_K_1_ treatments had significantly lower organic acid content than the other treatments. Therefore, balanced fertility can significantly reduce the content of organic acids in jujube fruit, and has a significant effect on changing the taste of jujube fruit. The sugar acids treated with N_3_P_1_K_3_ and N_3_P_3_K_1_ were 2.42% and 2.64% higher than the control, respectively. This has a lot to do with their high sugar content and low acid content. The total flavonoid content of K_3_ and N_1_P_3_K_3_ treatments was significantly higher than that of the other treatments. The total flavonoid content of P_2_K_2_, N_2_K_2_ and N_1_P_3_K_3_ treatments was also higher than that of the control, indicating that potassium fertilizer has a significant effect on increasing the total flavonoid content of pear jujube. (Table 3)

**Table 3.** Effect of N, P and K on the quality of jujube

| Treatment | Soluble solids (%) | Water content (%) | Total sugar (%) | Antitype Vitamin C (mg/100g) | Sugar- acid ratio | Total flavone (mg/100g) |
| --- | --- | --- | --- | --- | --- | --- |
| CK | 15.92±0.20 | 76.99±1.48 | 14.3±0.08 | 260.11±1.95 | 7.06±0.49 | 108.54±1.70 |
| $N_3$ | 16.02±0.39 | 79.29±1.20 | 16.2±0.23 | 258.57±1.68 | 11.78±1.29 | 117.04±0.72 |
| $P_3$ | 16.51±0.31 | 81.20±1.45 | 13.6±0.11 | 273.18±0.58 | 5.06±0.24 | 125.03±0.82 |
| $K_3$ | 17.12±0.06 | 81.33±0.25 | 13.8±0.14 | 285.76±1.13 | 5.61±0.06 | 138.21±1.70 |
| $P_2K_2$ | 18.11±0.30 | 78.87±1.78 | 16.0±0.20 | 249.56±0.73 | 10.38±0.27 | 120.90±1.10 |
| $N_2K_2$ | 17.18±0.11 | 80.33±1.09 | 15.3±0.16 | 268.23±4.19 | 9.01±0.29 | 111.31±1.45 |
| $N_2P_2$ | 16.98±0.05 | 80.74±2.14 | 16.4±0.19 | 262.91±2.29 | 9.83±0.51 | 80.59±1.81 |
| $N_1P_3K_3$ | 15.63±0.45 | 81.71±0.87 | 13.3±0.09 | 241.51±6.45 | 10.63±0.62 | 138.99±1.25 |
| $N_3P_1K_3$ | 16.65±0.06 | 80.46±0.55 | 13.5±0.14 | 264.80±1.68 | 17.12±1.03 | 125.66±1.26 |
| $N_3P_3K_1$ | 15.79±0.17 | 80.93±0.78 | 16.7±0.15 | 246.22±0.49 | 18.66±4.18 | 85.28±1.79 |

**Table 4.**
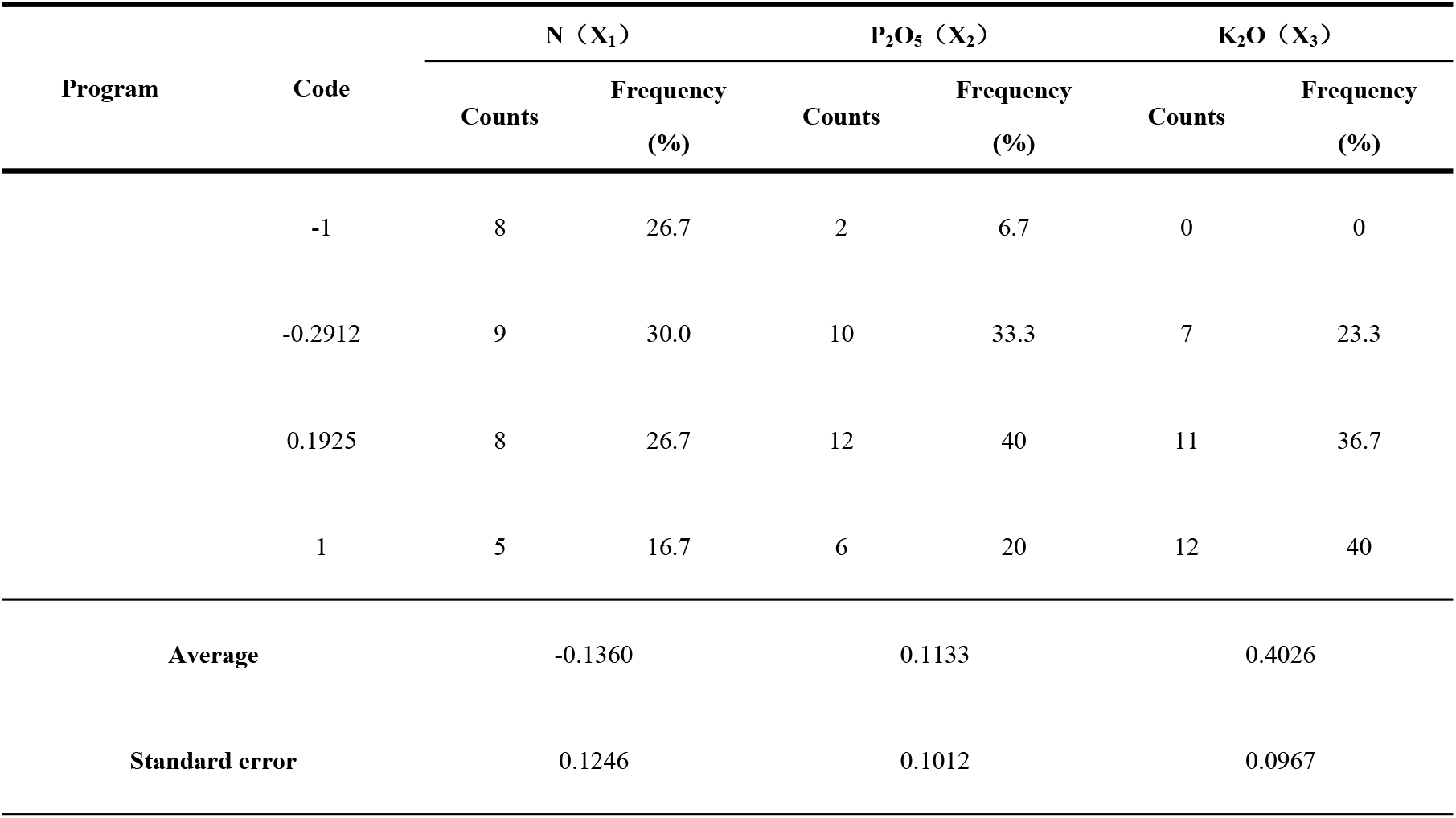
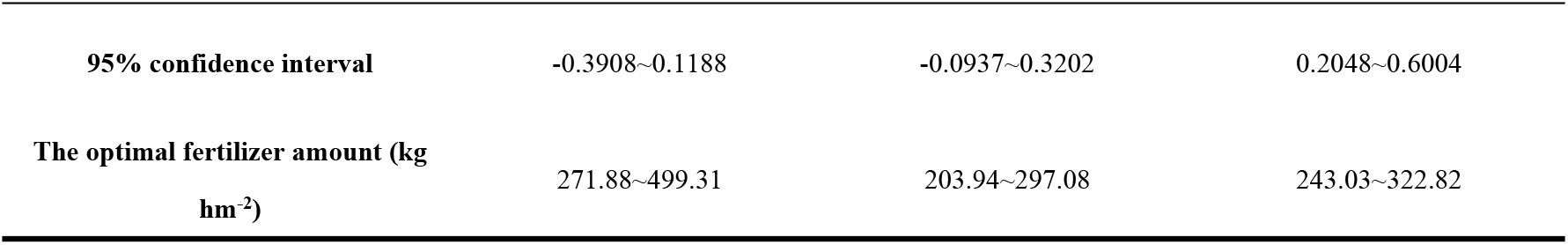
Options rates of N, P, K fertilizer for realizing the yield goat of 18000∼25000 kg hm^-2^

| Program | Code | N ($X_1$) | | P<sub>2</sub>O<sub>5</sub> ($X_2$) | | K<sub>2</sub>O ($X_3$) | |
| --- | --- | --- | --- | --- | --- | --- | --- |
|  |  | Counts | Frequency (%) | Counts | Frequency (%) | Counts | Frequency (%) |
|  | -1 | 8 | 26.7 | 2 | 6.7 | 0 | 0 |
|  | -0.2912 | 9 | 30.0 | 10 | 33.3 | 7 | 23.3 |
|  | 0.1925 | 8 | 26.7 | 12 | 40 | 11 | 36.7 |
|  | 1 | 5 | 16.7 | 6 | 20 | 12 | 40 |
| Average |  | -0.1360 |  | 0.1133 |  | 0.4026 |  |
| Standard error |  | 0.1246 |  | 0.1012 |  | 0.0967 |  |

|  |  |  |  |
| --- | --- | --- | --- |
| 95% confidence interval | -0.3908~0.1188 | -0.0937~0.3202 | 0.2048~0.6004 |
| The optimal fertilizer amount (kg<br>hm<sup>-2</sup>) | 271.88~499.31 | 203.94~297.08 | 243.03~322.82 |

### Establishment and Examination of Jujube Yield and Nitrogen, Phosphorus and Potassium Fertilization Models

The repeated data of the output of pear jujube in each treatment is shown in Table 2. The coding values of nitrogen, phosphorus and potassium fertilizers are independent variables and the output is dependent variable. In this way, the regression equation between red jujube yield and nitrogen, phosphorus, and potassium fertilizers was established. According to the output of pear jujube, the linear regression analysis of Microsoft Excel is used to obtain the code value effect functions of Y and N (X_1_), P (X_2_), and K (X_3_) of pear jujube in the region. Refer to Gong Jiang et al. (2011).

Y=24948.135-360.097X_1_+666.971X_2_+2070.209X_3_-387.043X_1_X_2_-1357.153X_1_X_3_+599.773X_2_ X_3_-1071.141X_1_^2^-1846.668X_2_^2^-1009.782X_3_^2^

F=101.483>F_0.01_(9, 20)=3.46 of the regression equation. The linear determination coefficient R^2^=0.9786, which reached a very significant level. This shows that the established regression equations for yield and fertilizer benefits are well simulated. Therefore, the model has a good prediction effect on the output of pear jujube.

### Analysis of Yield Effect Function

The main effect analysis of each fertilization element was carried out. Since the regression equations of nitrogen, phosphorus, and potassium fertilizers on yield have been replaced by dimensionless coding, the absolute values of the partial regression coefficients are directly compared. This can reflect the importance of each factor. From the first term of the regression model, the absolute values of the coefficients of nitrogen, phosphorus, and potassium are 360.097, 666.971, and 2070.209, respectively. It shows that potassium fertilizer has the largest effect on the jujube yield; phosphate fertilizer is the second; nitrogen fertilizer is the smallest. The partial regression coefficient of the first term of X_1_ is negative. It shows that the effect of nitrogen fertilizer on the output of pear jujube is negative. The partial regression coefficients of X_2_ and X_3_ are positive. Therefore, the effect of phosphate fertilizer and potassium fertilizer on the output of pear jujube is positive.

The dimensionality reduction method is used for the equation. It studies the single factor effects of nitrogen, phosphorus and potassium on the yield of pear jujube. Any two of the three independent variables in the model are fixed at zero level to obtain the single factor effect equation.

Effect of nitrogen fertilizer on yield: Y= 24948.135 -360.097X_1_ -1071.141X_1_^2^

Effect of phosphate fertilizer on yield: Y= 24948.135 + 666.971X_2_-1846.668X_2_^2^

Effect of potassium fertilizer on yield: Y= 24948.135 + 2070.209X_3_-1009.782X_3_^2^

The curve of the influence of nitrogen and phosphorus on the output of jujube is a parabola with an opening downward. When the amount of fertilization is small, nitrogen and phosphate fertilizer can obviously promote the yield of jujube. But after a certain amount of fertilization, the yield showed a downward trend with the increase of nitrogen and phosphate fertilizer. Within the range of fertilization used in this experiment, the effect of potassium fertilizer on the yield of pear jujube conforms to the law of diminishing returns of fertilizer benefits. That is, when the level of potassium application is low, the yield of pear jujube increases with the increase in the amount of potassium fertilizer application, but the increase rate continues to decrease.(Fig.1).

**Fig 1.**
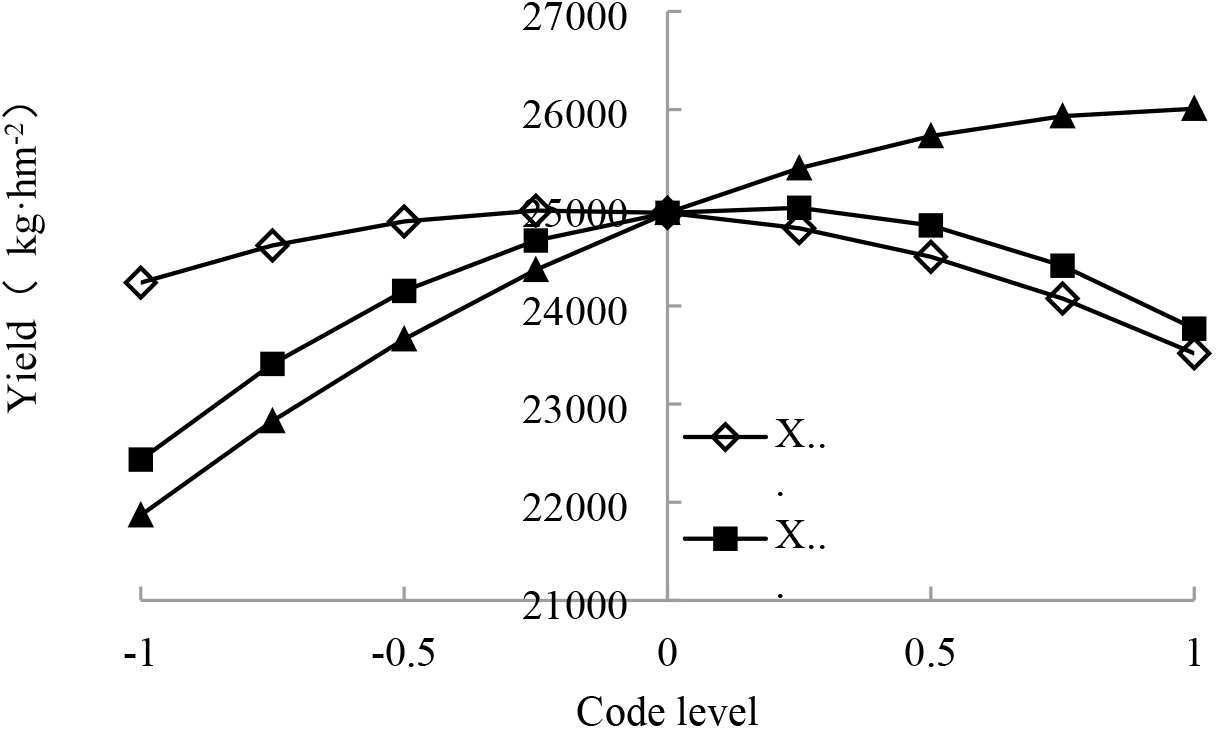
Analysis of single factor effects

### Optimize the fertilization mode according to the yield

According to the four levels of code values X_1_, X_2_ and X_3_, T = 4^3^ = 64 fertilization combinations are formed. The target yield is selected to be 23000 ∼ 27000 kg hm^-2^ (n = 30) for frequency analysis. Within this yield range, In this yield range, the optimal fertilizer application rates of nitrogen (N), phosphorus (P_2_O_5_) and potassium (K_2_O) are 271.88 ∼ 499.31 kg hm^-2^, 203.94 ∼ 297.08 kg hm^-2^, and 243.03 ∼ 322.82 kg hm^-2^. (Table 2)

### Comprehensive Quality Score of Red Jujube and Establishment and Test of Nitrogen, Phosphorus and Potassium Fertilization Models

Table 5 shows the data required for the comprehensive evaluation of the quality of red dates.

**Table 5.** Effect of N, P and K on the quality of jujube

| Treatment | Total flavone<br>(mg/100g) | Sugar- acid<br>ratio | Antitype<br>Vitamin C<br>(mg/100g) | Water content<br>(%) | Soluble solids<br>(%) | Fruit shape<br>index | Average value |
| --- | --- | --- | --- | --- | --- | --- | --- |
| CK | 108.54±1.70 | 7.06±0.49 | 260.11±1.95 | 76.99±1.48 | 15.92±0.20 | 0.841±0.050 | 76.64±1.30e |
| N<sub>3</sub> | 117.04±0.72 | 11.78±1.29 | 258.57±1.68 | 79.29±1.20 | 16.02±0.39 | 0.832±0.043 | 83.69±1.65bc |
| P<sub>3</sub> | 125.03±0.82 | 5.06±0.24 | 273.18±0.58 | 81.20±1.45 | 16.51±0.31 | 0.863±0.038 | 80.08±0.21d |
| **K<sub>3</sub>** | 138.21±1.70 | 5.61±0.06 | 285.76±1.13 | 81.33±0.25 | 17.12±0.06 | 0.908±0.014 | 85.26±0.52bc |
| **P<sub>2</sub>K<sub>2</sub>** | 120.90±1.10 | 10.38±0.27 | 249.56±0.73 | 78.87±1.78 | 18.11±0.30 | 0.879±0.048 | 84.00±1.15bc |
| **N<sub>2</sub>K<sub>2</sub>** | 111.31±1.45 | 9.01±0.29 | 268.23±4.19 | 80.33±1.09 | 17.18±0.11 | 0.919±0.025 | 81.85±0.66cd |
| **N<sub>2</sub>P<sub>2</sub>** | 80.59±1.81 | 9.83±0.51 | 262.91±2.29 | 80.74±2.14 | 16.98±0.05 | 0.888±0.016 | 75.30±1.41e |
| **N<sub>1</sub>P<sub>3</sub>K<sub>3</sub>** | 138.99±1.25 | 10.63±0.62 | 241.51±6.45 | 81.71±0.87 | 15.63±0.45 | 0.890±0.059 | 86.72±0.50b |
| **N<sub>3</sub>P<sub>1</sub>K<sub>3</sub>** | 125.66±1.26 | 17.12±1.03 | 264.80±1.68 | 80.46±0.55 | 16.65±0.06 | 0.874±0.061 | 92.67±1.53a |
| **N<sub>3</sub>P<sub>3</sub>K<sub>1</sub>** | 85.28±1.79 | 18.66±4.18 | 246.22±0.49 | 80.93±0.78 | 15.79±0.17 | 0.924±0.052 | 84.37±4.81bc |

The coding values of nitrogen, phosphorus and potassium fertilizers are independent variables and the comprehensive quality score is the dependent variable. In this way, a regression equation between the quality of red dates and nitrogen, phosphorus, and potassium fertilizers was established.

Y=79.957+2.133X_1_+0.448X_2_+5.135X_3_-1.743X_1_X_2_+0.354X_1_X_3_+0.475X_2_X_3_+4.797X_1_^2^+0.511 X_2_^2^+0.010X_3_^2^

F=21.95>F_0.01_(9, 20)=3.46 of the regression equation. The linear determination coefficient R^2^ = 0.8667, which reached a significant level. It shows that the established regression equation of the comprehensive quality fertilizer efficiency of red dates is a good simulation of the predicted value and the actual value. Therefore, the model has a good predictive effect on the quality of red dates.

### Analysis of quality effect function of pear-jujube

The main effect analysis of each fertilization element was carried out. Since the regression equations of nitrogen, phosphorus, and potassium fertilizers on yield have been replaced by dimensionless coding, the absolute values of the partial regression coefficients are directly compared. This can reflect the importance of each factor. The absolute values of the partial regression coefficients of X1, X2 and X3 in the regression model are 2.133, 0.448 and 5.135, respectively. It shows that under drip irrigation conditions, potassium fertilizer has the greatest influence on the quality of red dates. Nitrogen fertilizer followed. Phosphate fertilizer is the smallest. Each partial regression coefficient is positive. It shows that nitrogen, phosphorus and potassium have positive effects on the quality of red dates.

The dimensionality reduction method is used for the equation. It studies the single factor effects of nitrogen, phosphorus and potassium on the quality of pear jujube. Any two of the three independent variables in the model are fixed at zero level to obtain the single factor effect equation.

Effect of nitrogen fertilizer on quality: Y=79.957 + 2.133X_1_ + 4.797X_1_^2^

Effect of nitrogen fertilizer on quality: Y=79.957 + 0.448X_2_ + 0.511X_2_^2^

Effect of nitrogen fertilizer on quality: Y=79.957 + 5.135X_3_ + 0.010X_3_^2^

The effect curve of nitrogen fertilizer on the comprehensive score of pear jujube quality is a parabola with an upward opening. With the increase of fertilization, the comprehensive quality score of pear jujube gradually decreased. When the application amount of nitrogen fertilizer exceeds a certain fertilization level, the comprehensive quality score of t pear jujube will increase as the application rate of nitrogen fertilizer increases. Phosphate fertilizer has little effect on the comprehensive score of pear jujube quality. Potassium fertilizer is positively correlated with the comprehensive score of pear jujube. Within the fertilization range of the experimental design, the comprehensive quality score of red jujube increased with the increase of potassium fertilizer. (Fig.2)

**Fig 2.**
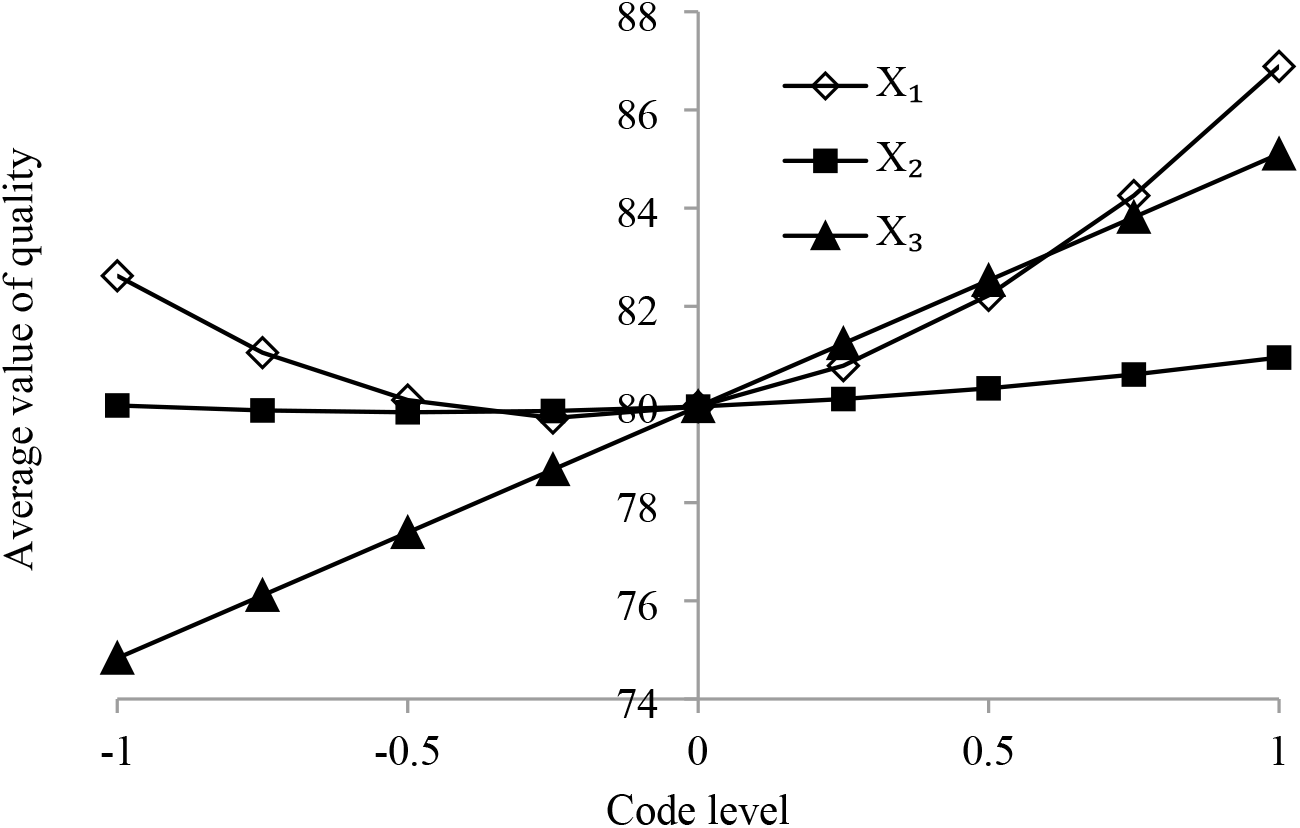
Analysis of single factor effects

### Optimizing fertilization mode based on quality

According to the four levels of code values X_1_, X_2_ and X_3_, T = 4^3^ = 64 fertilization combinations are formed. The comprehensive score of quality is selected to be above 85 points (n = 22) for frequency analysis. When the comprehensive score of quality is above 85 points, the optimized fertilization amount is nitrogen (N) 406.93 ∼ 736.69 kg hm^-2^, phosphorus (P_2_O_5_) 157.95 ∼ 306.39 kg hm^-2^, and potassium (K_2_O) 285.47 ∼ 376.32 kg hm^-2^. (Table 6)

**Table 6.** Frequency distribution of factors for quality average value over 85

| Program | Code | N (X<sub>1</sub>) |  | P<sub>2</sub>O<sub>5</sub> (X<sub>2</sub>) |  | K<sub>2</sub>O (X<sub>3</sub>) |  |
| --- | --- | --- | --- | --- | --- | --- | --- |
|  |  | Counts | Frequency (%) | Counts | Frequency (%) | Counts | Frequency (%) |
|  | -1 | 5 | 22.73 | 5 | 22.73 | 0 | 0 |
|  | -0.2912 | 2 | 9.09 | 5 | 22.73 | 3 | 13.64 |
|  | 0.1925 | 4 | 18.18 | 6 | 27.27 | 5 | 22.73 |
|  | 1 | 11 | 50 | 6 | 27.27 | 14 | 63.63 |
| Average |  | 0.2813 |  | 0.0318 |  | 0.6404 |  |
| Standard error |  | 0.1776 |  | 0.1586 |  | 0.1083 |  |
| 95% confidence interval |  | -0.0882~0.6507 |  | -0.2981~0.3616 |  | 0.4152~0.8656 |  |
| The optimal fertilizer amount(kg hm<sup>-2</sup>) |  | 406.93~736.69 |  | 157.95~306.39 |  | 285.47~376.32 |  |

## Discusses

The application of nitrogen, phosphorus and potassium fertilizers can obviously promote the yield and quality of pear jujube. Fertilization is beneficial to the growth of jujube fruit. Zhang et al [17] studied the seasonal dynamics of leaf nutrients in densely planted pear jujube. The contents of nitrogen, phosphorus and potassium in the leaves decreased rapidly during the fruit development period. The decreasing nutrient is transferred to the fruit for fruit growth. The results are consistent with this test. The effect of three fertilizers on the output of pear jujube is potassium fertilizer> phosphate fertilizer> nitrogen fertilizer. This is consistent with the research results of Chen et al. [16] and Chai [27] on gray jujube. In this experiment, single phosphate or potash fertilizer can effectively increase the water content of fruit. In the case of applying nitrogen fertilizer, adding potassium fertilizer or phosphate fertilizer alone or potassium and phosphate fertilizer can effectively increase the water content of jujube fruit. The mixed application of phosphate and potassium fertilizer has no significant effect on increasing the water content of jujube fruit. It shows that the application of fertilizer will increase the content of mineral elements in the soil. This is beneficial to the accumulation of mineral elements in plants. The increase of mineral elements in the cells will promote the cells to absorb water. Therefore, under drip irrigation, fertilization can increase the water content of jujube fruit. This is consistent with the research results. Different fertilization treatments can increase the water content in the fruit.

Chai [27] research shows that increasing the application of potassium fertilizer can increase the sugar and vitamin C content in jujube fruit. The Vc content of K_3_ was the highest in this experiment, which is consistent with Chai’s conclusion. But the highest total sugar content is N_3_P_3_K_1_ treatment. The sugar content of K_3_ treatment was lower than the control 0.29g/100g. The total sugar content of N_2_K_2_ and P_2_K_2_ treatments was not significantly different from N_3_P_3_K_1_ treatment. Analyze the possible reason for the negative effect caused by excessive potassium fertilizer. The specific mechanism needs further study. Lin et al [28] showed that the application of phosphorus and potassium fertilizers on the quality of citrus showed that single application of phosphorus, potassium fertilizer and combined application of phosphorus and potassium fertilizers can increase the content of soluble sugar in citrus. In this study, single application of phosphate fertilizer or potassium fertilizer had little effect on total sugar and reducing sugar content. Combined application of phosphorus and potassium can significantly increase the content of total sugar and reducing sugar. Analyze the reasons for the difference between the results of Lin [28] may be that the appropriate amount of potassium fertilizer is an important factor for increasing the sugar content in jujube fruit, and excessive potassium fertilizer will have a certain inhibitory effect. The specific mechanism needs to be further studied. Phosphate fertilizer and potassium fertilizer alone will increase the acid content in jujube fruit. Other treatments can reduce the content of organic acids in jujube fruit. High levels of phosphate and potassium fertilizers, and low levels of nitrogen fertilizer have a promoting effect on the total flavonoids content in jujubes. Therefore, proper fertilizer application should be paid attention to when applying fertilizers.

3-factor 2 times D-saturated optimal design has been successfully applied to Platycodon grandiflorus [30], Astragalns membranaceus [31], Onion [26] Research on optimal fertilization and cultivation conditions of other crops. Compared with the orthogonal combination design, the design requires less experimental processing. Model analysis can get the best test results, so it is widely used. In this experiment, the correlation analysis of the comprehensive score of red jujube yield and quality showed that the output and quality of red jujube were significantly correlated at the level of person 0.01. This is consistent with Zhao et al [29] about researching on tomato yield and quality. The results of this experiment showed that the application of nitrogen, phosphorus and potassium fertilizers had a significant effect on the yield and quality of Pear-jujube. Their increasing effect on yield is in order of potassium> phosphorus> nitrogen. Compared with Chai [27] and other studies on soil fertility of Xinjiang grey dates, the soils in loess hilly areas have higher contents of available nitrogen and available potassium, and lower contents of available phosphorus and organic matter. However, the sequence of nitrogen, phosphorus, and potassium fertilizers on the output of gray jujube is consistent with this study. It means that the jujube tree needs to apply suitable fertilizer to obtain high yield. The order of action on the quality of pear-jujube is potassium> nitrogen> phosphorus. Jujube trees are potassium-loving plants, and potash fertilizer application should be strengthened during planting. The optimization results of the fertilization model show that under the conditions of this experiment, when the target yield is 23000 ∼ 27000 kg hm^-2^, in the 95% confidence interval, the optimal fertilization amount is nitrogen (N) 272 ∼ 499 kg hm^-2^, phosphorus (P_2_O_5_) 204 ∼ 297 kg hm^-2^, potassium (K_2_O) 243 ∼ 323 kg hm^-2^, the demand of nitrogen fertilizer and potassium fertilizer in this result are lower than the results of Wang et al [32] research on Xinjiang Junzao. The demand for phosphate fertilizer is similar.

## Conclusions

When the target yield of pear jujube is in the range of 23000 ∼ 27000 kg hm^-2^, the optimized fertilization amount is nitrogen (N) 271.88 ∼ 499.31 kg hm^-2^, phosphorus (P_2_O_5_) 203.94 ∼ 297.08 kg hm^-2^, and potassium (K_2_O) 243.03 ∼ 322.82 kg hm^-2^.

When the comprehensive score of pear jujube quality is above 85 points, the optimized fertilization amount is nitrogen (N) 406.93 ∼ 736.69 kg hm^-2^, phosphorus (P_2_O_5_) 157.95 ∼ 306.39 kg hm^-2^, and potassium (K_2_O) 285.47 ∼ 376.32 kg hm^-2^.

Under the conditions of this experiment, the optimized fertilization rate for the target yield of red jujube is 23000 ∼ 27000 kg hm^-2^, and the comprehensive quality score is above 85 points is nitrogen (N) 406.93 ∼ 499.31 kg hm^-2^, phosphorus (P_2_O_5_) 203.94 ∼ 297.08 kg hm^-2^, Potassium (K_2_O) 285.47 ∼ 322.82 kg hm^-2^.

